# Reversion of antibiotic resistance in drug-resistant bacteria using non-steroidal anti-inflammatory drug benzydamine

**DOI:** 10.1101/2021.05.07.443075

**Authors:** Yuan Liu, Ziwen Tong, Jingru Shi, Tian Deng, Ruichao Li, Xia Xiao, Zhiqiang Wang

**Affiliations:** College of Veterinary Medicine, Yangzhou University, Yangzhou 225009, China; Institute of Comparative Medicine, Yangzhou University, Yangzhou 225009, China; Jiangsu Co-innovation Center for Prevention and Control of Important Animal Infectious Diseases and Zoonoses, Yangzhou University, Yangzhou 225009, China; Joint International Research Laboratory of Agriculture and Agri-Product Safety, the Ministry of Education of China, Yangzhou University, Yangzhou 225009, China

**Keywords:** antimicrobial resistance, antibiotic adjuvant, benzydamine, multidrug-resistant bacteria

## Abstract

Antimicrobial resistance has been a growing concern that gradually undermines our tradition treatment regimen. The fact that few antibacterial drugs with new scaffolds or targets have been approved in the past two decades aggravates this crisis. Repurposing previously approved drugs as potent antibiotic adjuvants offers a cost-effective strategy to mitigate the development of resistance and tackle the increasing infections by multidrug-resistant (MDR) bacteria. Herein, we found that benzydamine, a widely used non-steroidal anti-inflammatory drug in clinic, remarkably potentiated broad-spectrum antibiotic-tetracyclines activity against a panel of clinical important resistant pathogens, including MRSA, VRE, MCRPEC and *tet*(X)-positive Gram-negative bacteria. Further mechanistically experiments showed that benzydamine dissipated membrane potential (ΔΨ) in both Gram-positive and negative bacteria, which in turn upregulated the transmembrane proton gradient (ΔpH) and promoted the uptake of tetracyclines. Additionally, benzydamine exacerbated the oxidative stress by triggering the production of ROS and suppressing GAD system-mediated oxidative defensive. This mode of action explains the great bactericidal activity of the doxycycline-benzydamine combination against different metabolic states of bacteria including persister cells. As a proof-of-concept, the *in vivo* efficacy of this combination therapy was evidenced in multiple animal infection models. These findings revealed that benzydamine is a promising tetracycline antibiotics adjuvant and has the potential to address life-threatening infections by MDR bacteria.

## 1. Introduction

The prevalence of chromosome or plasmid-conferred resistance determinants have severely impair the efficacy of clinically available antibiotics, rendering the onset of the global antimicrobial resistance crisis (Harrison and Brockhurst, 2012). Among these pathogenic bacteria, of particular concerns are ESKAPE (Enterococcus, *Staphylococcus aureus, Klebsiella pneumoniae, Acinetobacter baumannii, Pseudomonas aeruginosa*, and *Enterobacter* species), which are responsible for the majority of nosocomial infections worldwide with high morbidity and mortality (De Oliveira et al., 2020; Ma et al., 2020). As the increasing incidence of drug resistance, including multidrug resistance and pandrug resistance (PDR), in these ESKAPE clinical isolates, bacterial infection associated diseases are becoming harder to treat. Notably, carbapenems, colistin and tigecycline are recognized as extremely crucial antibiotics and last-options against these drug-resistant bacteria.

However, the emergence of carbapenemase (Gupta et al., 2011), *mcr-1*-encoded phosphoethanolamine transferase (Liu et al., 2016) and *tet*(X)-mediated flavin-dependent (FAD) monooxygenase (He et al., 2019; Sun et al., 2019) in bacteria from animal and humans source completely extinguished our last hope. Meanwhile, few novel antibiotics entities with distinct scaffolds or modes of action have been approved for clinical use during the past decades due to the huge scientific and commercial challenges in the development of new drugs (Lewis, 2020; Liu et al., 2019a). There is a dire need to identify alternative strategies to address these infections.

Repurposing previously approved drugs as potential antibiotic adjuvants to reverse antibiotic resistance and restore antibiotic activity represents a simple but effective approach to counter this problem (Liu et al., 2019b; Wright, 2016). For example, our previous studies have showed that hypoglycemic drugs metformin could resensitive *tet*(A)-positive bacteria to tetracycline through disrupting the functions of efflux pumps (Liu et al., 2020b). Melatonin, which has been applied for treating sleep disturbances and circadian disorders, potentiated colistin activity against MCR-positive bacteria by enhancing the membrane damage (Liu et al., 2020c). Anti-HIV agent azidothymidine decreased Tet(X3/X4)-mediated bacterial resistance to tigecycline in *Escherichia coli* through specifically inhibiting DNA synthesis and suppressing resistance enzyme activity (Liu et al., 2020a; Zhou et al., 2020). Benzydamine is a locally-acting nonsteroidal anti-inflammatory drug with local anaesthetic and analgesic properties by selectively binding to prostaglandin synthetase (Avvisati et al., 2018; Nettis et al., 2002). Recently, benzydamine was found to inhibit osteoclast differentiation and bone resorption by down-regulating the expression of interleukin-1β (Son et al., 2020). In addition, benzydamine significantly reduced oral mucositis even at doses >50 Gy in head and neck cancer patients (Rastogi et al., 2017). However, the adjuvant potential of benzydamine to existing antibiotics is still unclear. In the present study, we characterized the synergistic activity of benzydamine with different classes of antibiotics, and found that it drastically potentiated tetracyclines activity against various MDR pathogens. Importantly, benzydamine dissipated membrane potential (ΔΨ) in both Gram-positive and negative bacteria, which in turn upregulated the transmembrane proton gradient (ΔpH) and promoted the uptake of tetracyclines. Meanwhile, benzydamine synergized with doxycycline on killing a spectrum of bacterial pathogens carrying *mec*A, *bla*_MBL_ and/or *mcr* genes, as well as *tet*(X) by triggering oxidative damage. Notably, benzydamine potently restored the doxycycline activity in multiple animal infection models infected by MDR MRSA T44 or *E. coli* B2. This study firstly revealed the therapeutic potential of benzydamine as a novel antibiotic adjuvant for the treatment of infection caused by MDR pathogens.

## 2. Results

### 2.1 Benzydamine potentiates doxycycline activity in both drug-susceptible and resistant bacteria

We first evaluated the synergistic activity of benzydamine with eight classes of antibiotics against multidrug-resistant bacteria *E. coli* B2 using checkerboard broth microdilution assays. Out of these drugs, colistin, ciprofloxacin and doxycycline showed synergistic activity with benzydamine, whereas kanamycin displayed an antagonistic effect with benzydamine **(Figure 1-figure supplement 1 and Table 1)**. Remarkably, the combination of benzydamine and doxycycline possessed the highest synergistic effect (FICI = 0.188), which enabled the MIC value of doxycycline decreased from 32 μg/mL to 2 μg/mL (16-fold). We further tested the potentiation of benzydamine to other tetracyclines, including tetracycline, oxytetracycline, minocycline and tigecycline. As expected, their antibacterial activity were all significantly improved in the presence of benzydamine **(Table 1)**. Subsequently, the checkerboard broth microdilution assays were applied to both sensitive and resistant bacteria. Interestingly, the combination of benzydamine and doxycycline showed synergy effect in all test bacteria, including hard-to-treat pathogenic bacteria methicillin-resistant *Staphylococcus aureus* (MRSA) T144 (FICI = 0.188), VRE A4 (FICI = 0.375), *bla*_NDM-5_-positive *E. coli* G6 (FICI = 0.375), *mcr-1*-carrying *K. pneumoniae* D120 (FICI = 0.375) and *tet*(X6)-positive *A. baumannii* C222 (FICI = 0. 5). Notably, this combination displayed a higher synergistic effect in drug-resistant bacteria than sensitive bacteria, suggesting that its activity is also related to the inhibition of resistance determinants **(Figure 1 and Table 2)**.

**Table 1.** Synergistic activity of benzydamine and antibiotics against MDR *E. coli* B2.

| Antibiotics | MIC<sup>a</sup> (μg/mL) | FIC index | MIC<sup>b</sup> (μg/mL) | Potentiation (fold)<sup>c</sup> |
| --- | --- | --- | --- | --- |
| Ampicillin | >128 | 2.0 | >128 | — |
| Meropenem | 64 | 2.0 | 64 | — |
| Rifampicin | 128 | 1.5 | 64 | 2 |
| Vancomycin | 128 | 2.0 | 128 | — |
| Kanamycin | 256 | >2 | >256 | — |
| Colistin | 8 | 0.5 | 2 | 4 |
| Ciprofloxacin | 16 | 0.375 | 4 | 4 |
| Doxycycline | 32 | 0.188 | 2 | 16 |
| Tetracycline | 128 | 0.25 | 16 | 8 |
| Oxytetracycline | 256 | 0.375 | 64 | 4 |
| Minocycline | 16 | 0.188 | 1 | 16 |
| Tigecycline | 2 | 1.0 | 1 | 2 |

<sup>a/b</sup>MICs of antibiotic in the absence or presence of 0.25×MIC of benzydamine.
<sup>c</sup>Degree of antibiotic potentiation in the presence of 0.25×MIC of benzydamine.
—, none of potentiation activity.

**Table 2.** Synergistic activity of benzydamine and doxycycline against drug-sensitive or resistant bacteria.

| Pathogens | MIC<sup>a</sup><br>(µg/mL) | FIC index | MIC<sup>b</sup><br>(µg/mL) | Potentialiation<br>(fold)<sup>c</sup> |
| --- | --- | --- | --- | --- |
| **Sensitive bacteria** |  |  |  |  |
| *S. aureus* ATCC 29213 | 0.5 | 0.5 | 0.125 | 4 |
| *E. coli* ATCC 25922 | 1 | 0.5 | 0.25 | 4 |
| *S. enteritidis* ATCC 13076 | 2 | 0.5 | 0.5 | 4 |
| *E. coli* MG1655 | 2 | 0.5 | 0.5 | 4 |
| **Resistant bacteria** |  |  |  |  |
| MRSA T144 | 16 | 0.188 | 1 | 16 |
| VRE A4 | 32 | 0.375 | 8 | 4 |
| *E. coli* G6 | 16 | 0.375 | 2 | 8 |
| *K. pneumoniae* D120 | 32 | 0.375 | 4 | 8 |
| *A. baumannii* C222 | 16 | 0.5 | 4 | 4 |

<sup>a/b</sup>MICs of antibiotic in the absence or presence of 0.25×MIC of benzydamine. <sup>c</sup>Degree of antibiotic potentiation in the presence of 0.25×MIC of benzydamine. —, none of potentiation activity.

**Figure 1.**
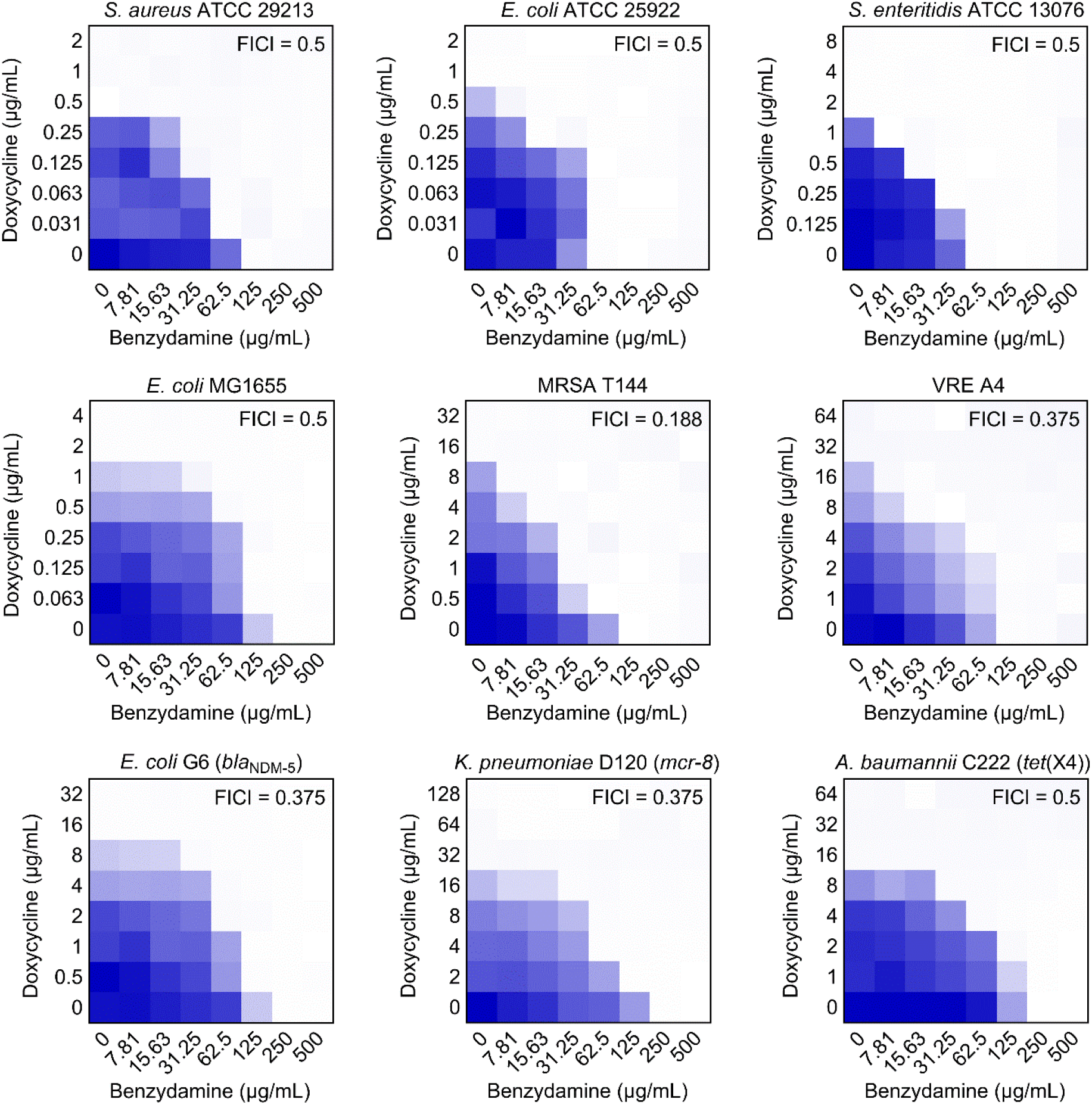
Synergistic activity between benzydamine and doxycycline against drug-sensitive and resistant bacteria by checkerboard assay, related to Table S3. Dark blue regions represent higher cell density. Data represent the mean OD_600_ nm of two biological replicates. Synergy is defined with FIC index ≤ 0.5

Next, we assessed whether the synergistic activity of this combination would result in increasing toxicity, including hemolytic activity on mammals RBCs and *in vivo* toxicity in mice (Lakshmaiah Narayana et al., 2020). Surprisingly, no detectable toxicity in hemolysis rate, body weight and blood biochemical analysis were found in the benzydamine-doxycycline combination treatment **(Figure 1-figure supplement 2 and 3)**. These data suggested that benzydamine was a safe and potent antibiotic adjuvant to tetracyclines.

### 2.2 Benzydamine dissipates the electric potential (ΔΨ) component of proton motive force and promotes the uptake of doxycycline

Our prior results have shown that benzydamine is a universal adjuvant to tetracycline antibiotics in all tested strains, but antagonize kanamycin activity in *E. coli* B2. We next assessed the interaction of benzydamine and kanamycin in a panel of bacteria. As a consequence, a significant antagonism effect were found in all bacteria (FICI > 2.0, **Figure 1 and Figure 1-figure supplement 4)**. The opposite action of benzydamine in combination with doxycycline or kanamycin inspired us to speculate on the mechanism of action of benzydamine may be directly related to the destruction of the bacterial proton motive force (PMF) (Farha et al., 2018). In bacteria, the transmembrane transfer of proton H^+^ by the respiratory chain results in an electrochemical gradient, named PMF. It consists of two parts, electric potential (ΔΨ) and pH difference (ΔpH) (Mitchell, 2011). Damage to one will be compensated by increasing another to achieve dynamic balance (Chen et al., 2008). Previous studies have indicated that the uptake of tetracyclines by bacterial cells depends on ΔpH, whereas aminoglycosides utilizes the ΔΨ component for transport, therefore, we concerned that benzydamine might target the ΔΨ component of PMF. To test our hypothesis, a fluorescent probe 3,3-dipropylthiadicarbocyanine iodide (DiSC_3_(5)) (Liu et al., 2020d) was used to assess membrane potential changes induced by doxycycline, benzydamine alone or their combination. After treatment of four representative strains (*S. aureus* ATCC 29213, MRSA T144, *E. coli* ATCC 25922 and *E. coli* B2; two Gram-positive and two Gram-negative bacteria; also two doxycycline-sensitive and two doxycycline-resistance bacteria) with 4-fold MIC of doxycycline, the fluorescence of Gram-positive bacteria hardly changed and Gram-negative bacteria slightly increased. However, treatment with 125-1,000 μg/mL of benzydamine resulted in rapid disruption of electric potential in a dose-dependent manner **(Figure 2A)**. Next, we measured the fluorescence with 1 to 4-fold MIC of doxycycline or combination with 250 μg/mL of benzydamine. The combination of benzydamine and doxycycline indeed resulted in increased fluorescence **(Figure 2B)**, suggesting that benzydamine is definitely a potential dissipator of ΔΨ. The extracellular pH values are also related to the PMF. A previous study demonstrated that the antibacterial activity of the dissipater of ΔΨ will be strengthened when the extracellular pH changed to the alkaline values (Farha et al., 2013). Consistent with the membrane potential results, the MIC of benzydamine were reduced by 8-fold due to the pH from 5.5 to 9.5 in the Gram-negative bacteria, and 4-fold change for Gram-positive bacteria **(Figure 2C)**. An intact PMF is required for the bacterial function in flagellar secretion, thus we next examined the integrity of PMF through swimming motility experiments (Brunelle et al., 2017). Exposure of four strains to sub-inhibitory concentrations of benzydamine drastically decreased bacterial motility **(Figure 2D)**, suggesting the impaired PMF in benzydamine-treated bacterial cells. These evidences demonstrated that benzydamine disrupted the PMF by targeting ΔΨ component.

**Figure 2.**
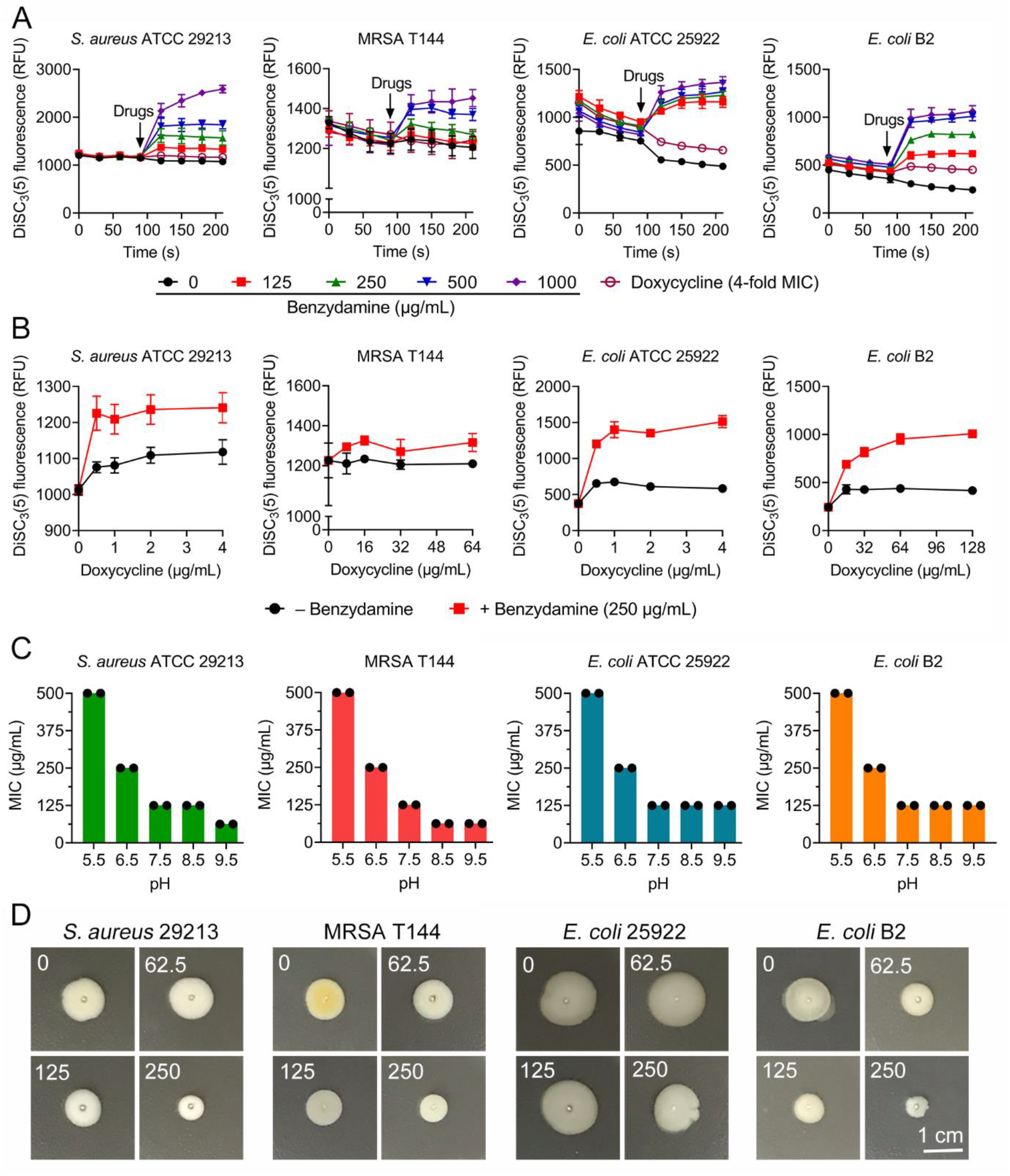
Benzydamine disrupts proton motive force (PMF) in both Gram-positive and Gram-negative bacteria. **(A)** Benzydamine dissipates membrane potential in bacteria. Fluorescence intensity of DiSC_3_(5) in *S. aureus* ATCC 29213, MRSA T144, *E. coli* ATCC 25922 and *E. coli* B2 after treatment with increasing concentrations of benzylamine and doxycycline (4-fold MIC) was monitored. Drugs were added into DiSC_3_(5)-probed cells at 90 s. **(B)** Combination of doxycycline and benzydamine (250 μg/mL) displays increased disruption on membrane potential compared with doxycycline alone. **(C)** Decreased MIC values of benzydamine against four bacteria in alkaline environment. ΔΨ becomes the main component of PMF as the external pH is shifted to an alkaline environment. **(D)** Benzydamine inhibits swimming motility of four bacterial strains. Overnight cultures were standardized to OD_600_ nm of 0.5, and inoculated on 0.3% agar plates for 48 h at 37°C. Scar bar, 1 cm.

There is a compensation mechanism that damage to one will be compensated by increasing another to maintain the dynamic balance of PMF. We have observed that benzydamine selectively disrupted the ΔΨ in before, so we further set out to test whether ΔpH will be compensatory upregulated. A membrane-permeable fluorescent probe termed BCECF-AM (Ozkan and Mutharasan, 2002) was used to monitor intracellular pH changes in four strains. Interestingly, completely opposite pH changes were observed in Gram-positive and negative bacteria. Benzydamine led to the acidification of the cytoplasm in G^+^, but alkalization of cytoplasm in G- **(Figure 3A)**. Nevertheless, both these actions triggered the upregulation of ΔpH in bacteria. Given that the increasing ΔpH by benzydamine may promote the uptake of tetracyclines, thus we determined the intracellular doxycycline accumulation after exposure to varying concentrations of benzydamine (Ejim et al., 2011). As expected, benzydamine supplementation remarkably enhanced the content of doxycycline in bacteria **(Figure 3B)**. Tetracyclines exhibit activity by specifically binding to the 30S subunit of the ribosome, thus inhibiting bacterial protein synthesis (Chopra and Roberts, 2001). Therefore, the uptake and accumulation of tetracyclines is of importance for its antibacterial activity. Collectively, these results suggested that benzydamine dissipated the ΔΨ, in turn upregulate the ΔpH, thereby promoted the uptake of tetracyclines.

**Figure 3.**
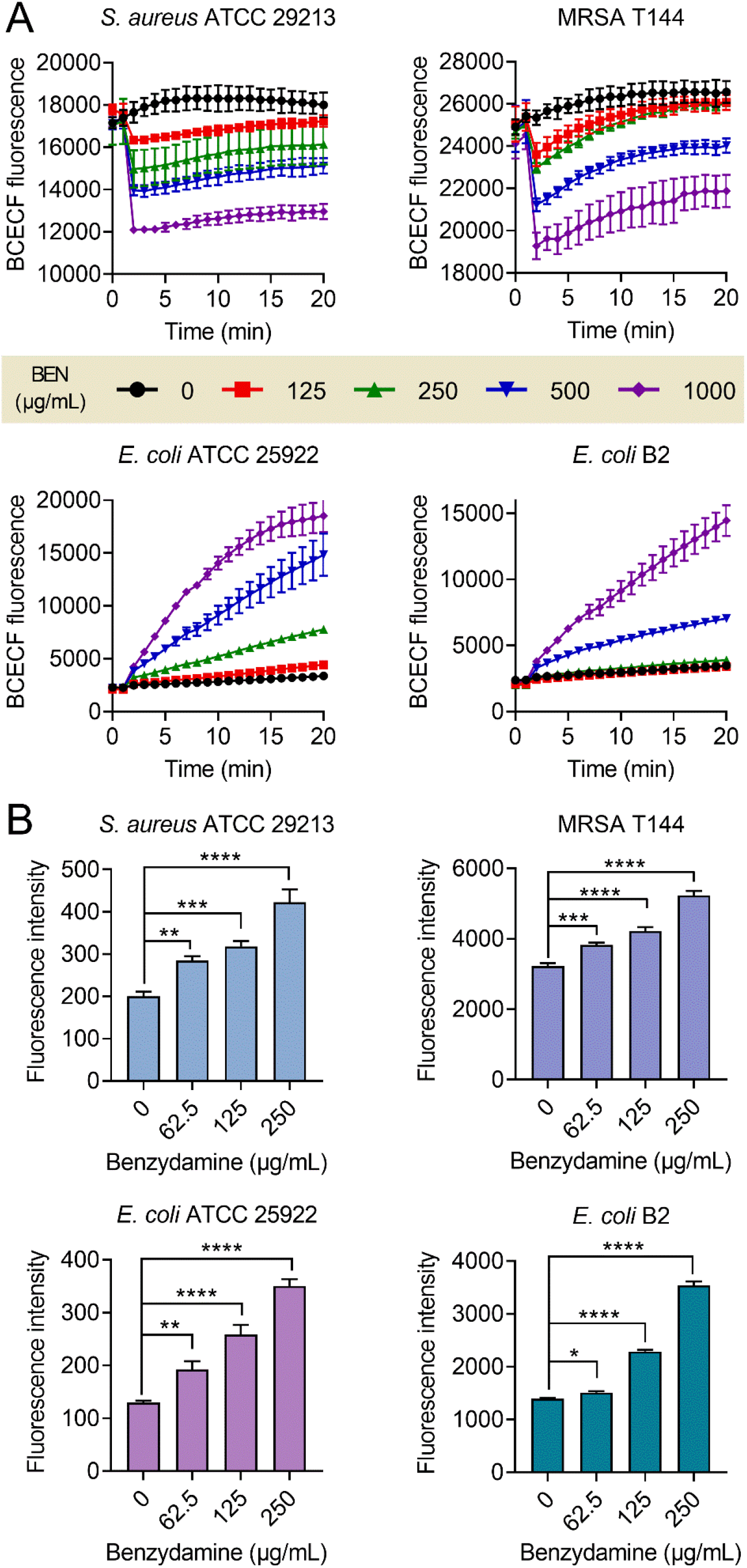
Benzydamine upregulates ΔpH and promotes the intracellular accumulation of doxycycline. **(A)** Upregulation of ΔpH in BCECF-AM-labeled bacterial cells after exposure to varying concentrations of benzydamine. In Gram-positive bacteria, benzydamine decreases fluorescence and the cytoplasmic pH. In contrast, benzydamine increases fluorescence and the cytoplasmic pH in Gram-negative bacteria (*E. coli* ATCC 25922 and *E. coli* B2). **(B)** Benzydamine supplementation dose-dependent promotes the intracellular accumulation of doxycycline in bacteria. Intracellular antibiotic content was determined by monitoring the fluorescence of doxycycline (excitation wavelength, 405 nm; emission wavelength, 535 nm). All data were presented as mean ± SD, and the significance was determined by non-parametric one-way ANOVA (\**P* < 0.05, \*\**P* < 0.01, ***P < 0.001, \*\*\*\**P* < 0.0001).

### 2.3 Doxycycline plus benzydamine is bactericidal against MDR bacteria and biofilm-producing bacteria

It has been widely acknowledged that tetracyclines belong to bacteriostatic antibiotics. We reasoned whether the benzydamine-doxycycline combination would possess bactericidal activity, which would markedly extend its therapeutic potential. To test this hypothesis, we performed time-killing experiments on various MDR pathogens. Impressively, a direct synergistic bactericidal effect was observed in rich growth conditions **(Figure 4A)**. Specifically, either 32 μg/mL doxycycline or 250 μg/mL benzydamine showed slight bactericidal activities. In comparison, the combination of doxycycline plus benzydamine (32 + 250 μg/mL) exhibited excellent bactericidal activity, especially for *E. coli* B2. Besides, to determine whether benzydamine has potency to combat metabolically repressed and non-replicating cells, we test the bactericidal activity of this combination in nutrient-free buffer. Remarkably, this combination retained potent bactericidal activity **(Figure 4B)**.

**Figure 4.**
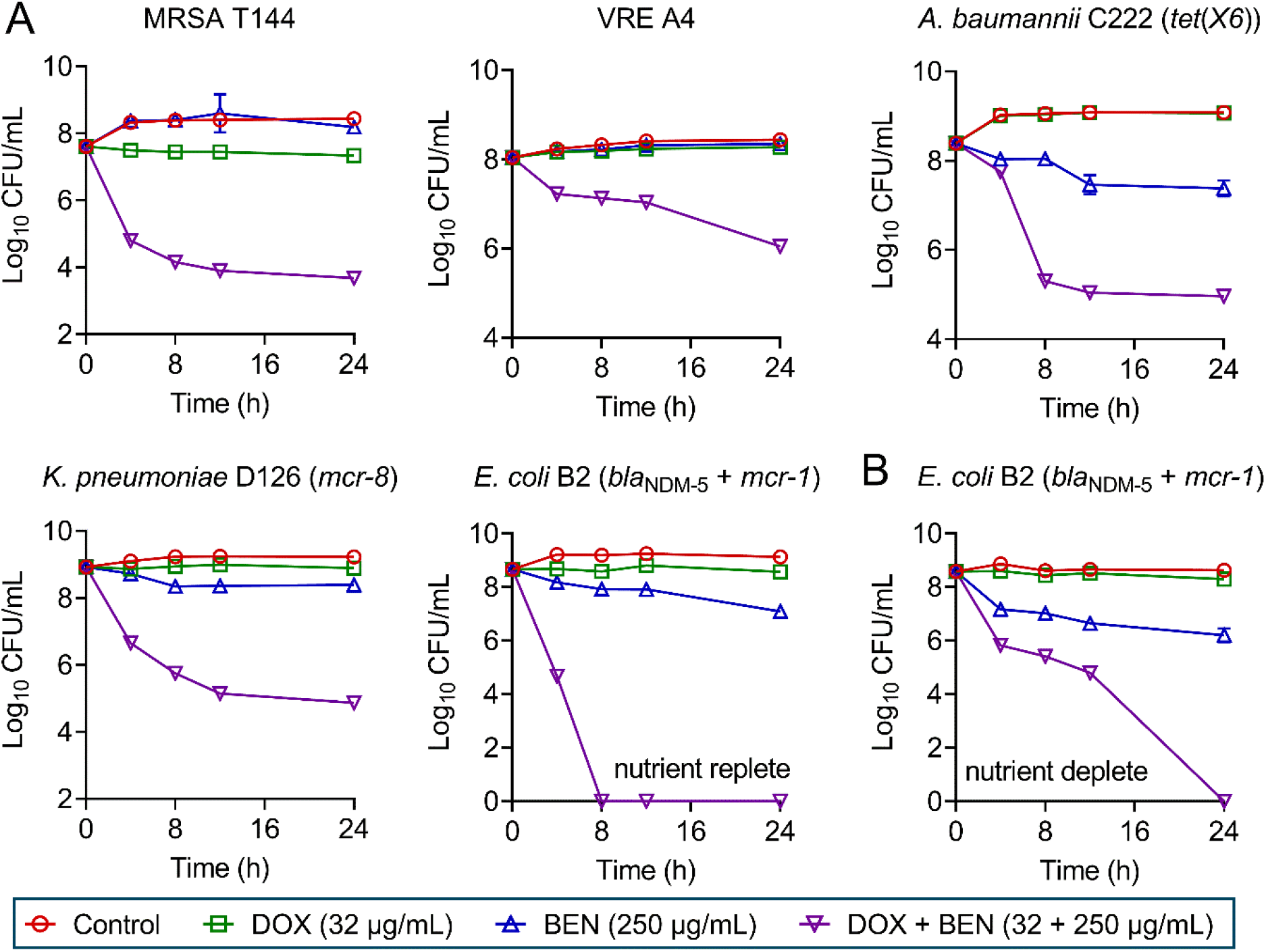
The combination of doxycycline and benzydamine is bactericidal against various drug-resistant bacteria. **(A)** Killing activity of doxycycline plus benzydamine in LB media against multidrug-resistant Gram-positive bacteria (MRSA T144 and VRE A4) and Gram-negative bacteria (*A. baumannii* C222, *K. pneumoniae* D126 and *E. coli* B2). **(B)** Killing activity of doxycycline plus benzydamine in PBS against *E. coli* B2. The initial cell density is about 10^8^ CFU/mL. All data were performed from three biological replicates and shown as mean ± SD.

The formation of antibiotic-tolerant biofilms greatly affect the efficacy of antibiotics (Hall and Mah, 2017; Yan and Bassler, 2019). To explore whether benzydamine supplementation can enhance doxycycline activity against the biofilm-producing bacteria, we performed the formation and eradication of biofilms experiments in the presence or absence of different concentrations of benzydamine. As shown in **Figure 4-figure supplement 1A**, the combination of benzydamine at 50 μg/mL with sub-MIC of doxycycline significantly inhibited biofilm formation of MRSA T144 and *E. coli* B2. Notably, in the biofilm inhibition assay, the benzydamine plus doxycycline at concentrations of ≤ 2 μg/mL did not have bacteriostatic activity against two test strains, indicating that the inhibition of biofilm formation at these concentrations was not due to the effect on bacterial growth. Besides, in the presence of benzydamine, the eradication effect of doxycycline on mature biofilm is significantly enhanced compared with doxycycline alone **(Figure 4-figure supplement 1B)**. Taken together, we unexpectedly found that the combination of doxycycline plus benzydamine displayed great bactericidal activity against various MDR pathogens in different metabolic states, including metabolically active cells, antibiotic-tolerant cells and biofilm-producing bacteria.

### 2.4 Benzydamine aggravates oxidative damage and inhibits the function of MDR efflux pumps

Having shown that the synergistic bactericidal activity of benzydamin-doxycycline combination, we reasoned that benzydamine may trigger other unknown modes of action except for the promotion of doxycycline uptake. To explore the underlying mechanisms, we performed transcription analysis of *E. coli* B2 under treatment with doxycycline or doxycycline plus benzydamine for 4 h. The comparison of treatment with combination to antibiotic alone revealed an up-regulation of 35 differentially expressed genes (DEGs) and down-regulation of 14 DEGs **(Figure 5-figure supplement 1A)**. Go annotation analysis showed that these DEGs were involved in biological processes, cellular components and molecular functions **(Figure 5-figure supplement 1B)**. KEGG enrichment analysis displayed that these DEGs with increased expression were involved in ribosome synthesis, and DEGs with repressed expression in glutamate metabolism and GABA shunt **(Figure 5-figure supplement 1C and D)**. Notably, genes with 30S and 50S subunit were up-regulated, which may be caused by increased accumulation of doxycycline that inhibits protein synthesis **(Figure 5A and source data 2)**. In contrast, multidrug efflux pumps related genes, glutamate decarboxylase (GAD) system associated genes, and acid resistance related gene was obviously decreased **(Figure 5B and source data 2)**.

**Figure 5.**
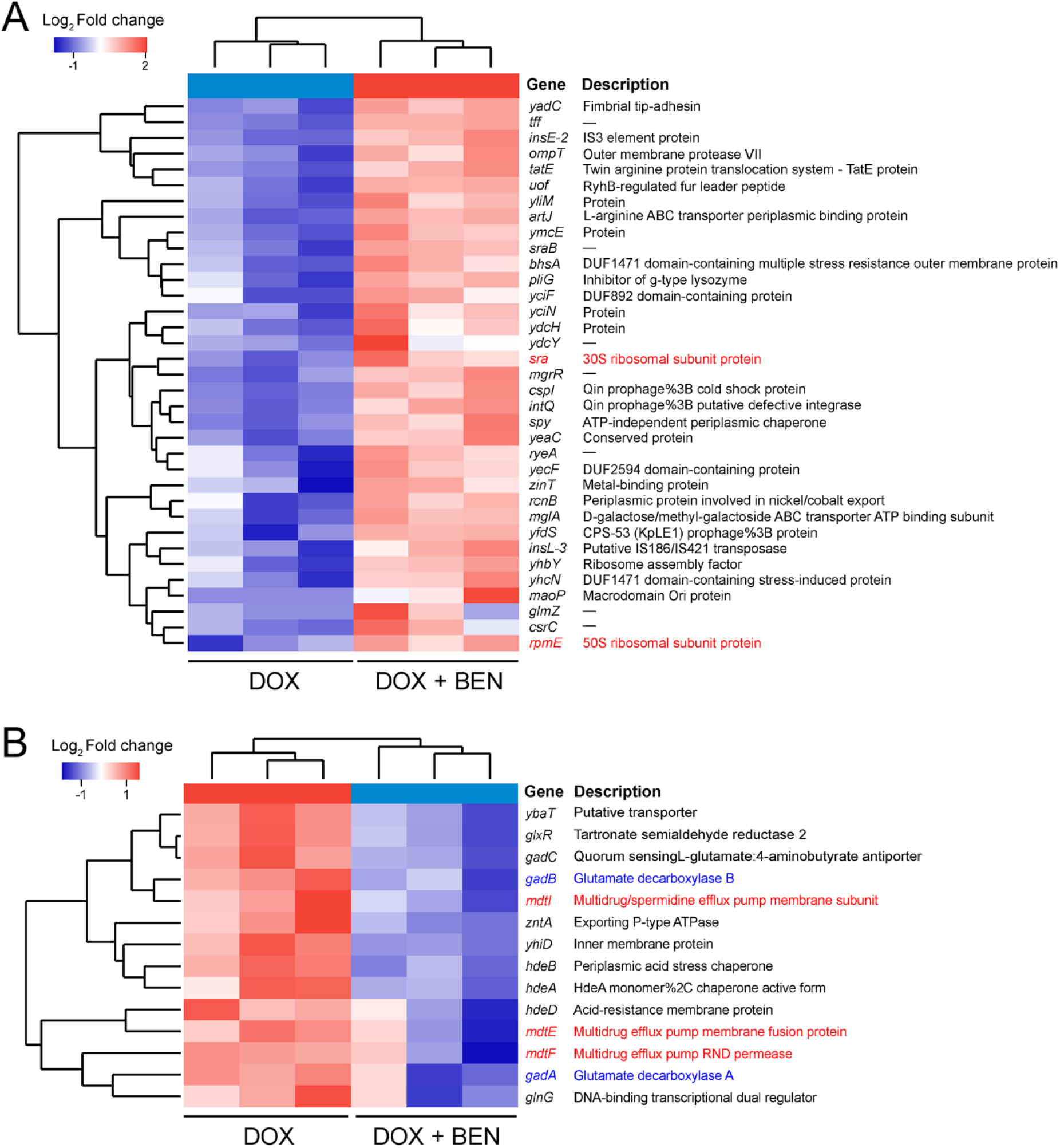
Differentially expressed genes (DEGs) of *E. coli* B2 after treatment with doxycycline plus benzydamine in comparison to doxycycline alone. Significant up-regulated **(A**, *P* < 0.05, Log2Fold change ≥ 1) and down-regulated DEGs **(B**, *P* < 0.05, Log2Fold change ≤ -1) in combination treatment group compared with doxycycline monotreatment.

Based on the transcription results, we next performed a series of phenotype experiments to elucidate the other functions of benzydamine. Firstly, we evaluated the efficacy of combination in the different pH growth environment via time-killing experiments. We found that it has almost no effect in the acid media, however, the combination of doxycycline with benzydamine showed manifest bactericidal activity while a change in pH to alkaline values. Under treatment 4 hours, the bacteria were all killed in the MHB broth at pH 8.5 and 9.5 **(Figure 6A)**. These data were in agreement with previous observation that antibacterial activity of benzydamine was strengthened in the alkaline conditions and the genes associated with acid resistance were down-regulated. Recently, several studies reported that GAD systems, which was downregulated in combination group, plays a critical role in protecting bacteria against oxidative stress (Boura et al., 2020; Feehily and Karatzas, 2013), thus we hypothesized that the potentiation of benzydamine to antibiotics may also correlate to enhanced oxidative damage. Thus, we tested the generation of reactive oxygen species (ROS) (Voorhees, 2003) in *E. coli* B2 after treatment with either benzydamine or in combination with doxycycline. Surprisingly, benzydamine markedly promoted the generation of ROS in a dose-dependent manner **(Figure 6B)**. Meanwhile, the combination treatment showed higher ROS levels compared to doxycycline monotreatment **(Figure 6C)**. Accordingly, ROS has been recognized as one of common mechanisms in antibiotic-mediated killing of bacteria. The over-production of ROS in benzydamine-doxycycline combination give an interpretation on their synergistic bactericidal activity. To further verify it, *N*-acetyl-*L*-cysteine (NAC), a ROS scavenger, was added in time-killing assays. As shown in **Figure 6D**, the potentiation of benzydamine to doxycycline was greatly impaired when incubation with 2 or 4 mM NAC. Finally, we used a fluorescent dye Rhodamine B to assay the function of efflux pump in bacteria under the treatment of benzydamine **(Figure 6E)**. As a result, it showed that the activity of efflux pump was significantly suppressed in the presence of 250-1,000 μg/mLbenzydamine, which further promoted the accumulation of doxycycline in the MDR bacteria. Collectively, these phenomenon together demonstrated that benzydamine enhanced oxidative damage through triggering the production of ROS and inhibiting the function of MDR efflux pumps **(Figure 6F)**.

**Figure 6.**
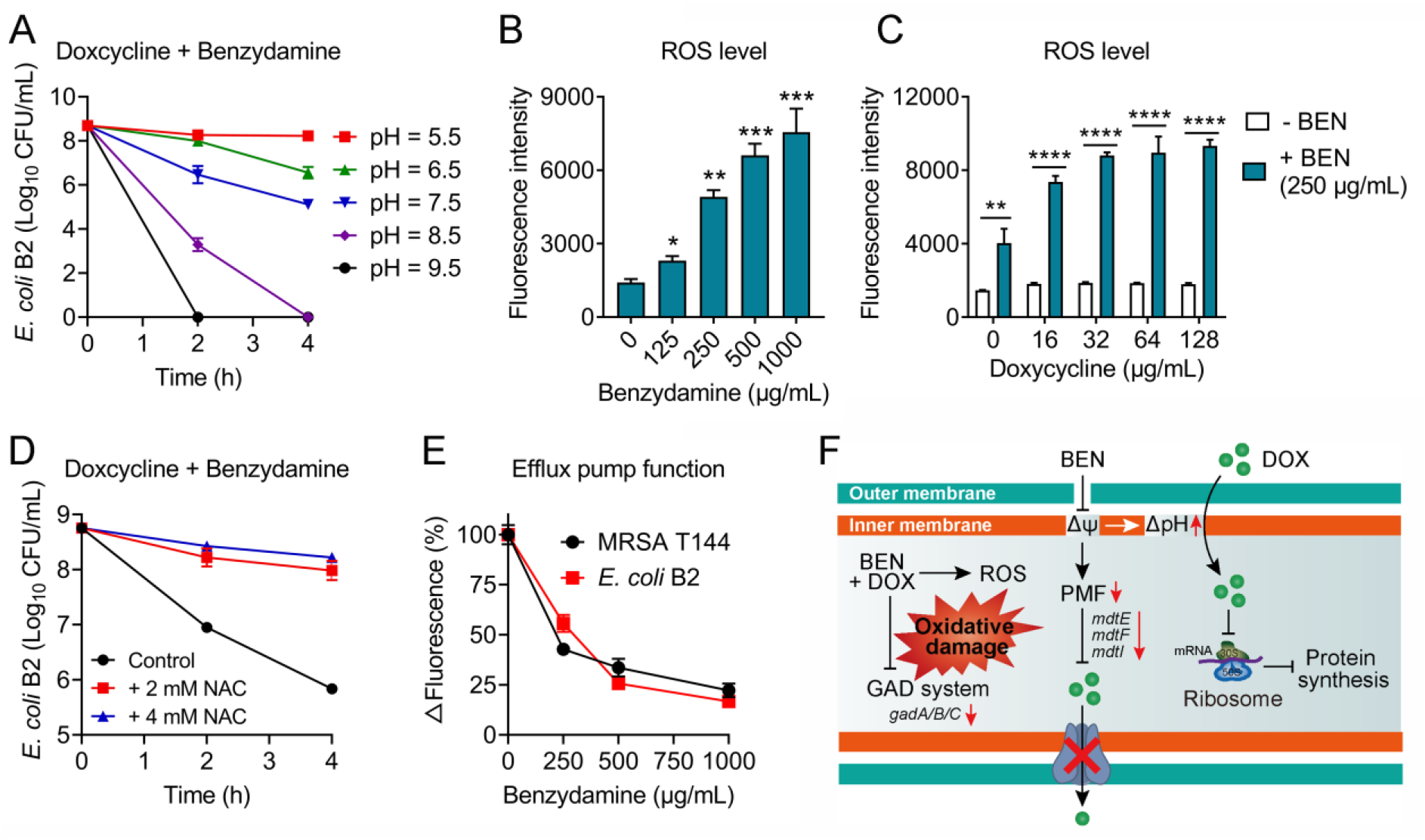
Benzydamine promotes oxidative damage and inhibits the functions of efflux pump in *E. coli*. **(A)** Time-killing curves of *E. coli* B2 after treatment with the combination of doxycycline and benzydamine in different pH media from 5.5 to 9.5. **(B)** Benzydamine promotes the production of ROS in a dose-dependent manner. **(C)** Doxycycline plus benzydamine (250 μg/mL) showed higher ROS generation compared to doxycycline alone. Fluorescence probe 2′,7′-dichlorodihydrofluorescein diacetate (DCFH-DA) was used to monitor the levels of ROS in cells (λexcitation = 488 nm, λemission = 525 nm). All data were presented as mean ± SD, and the significance was determined by non-parametric one-way ANOVA (\**P* < 0.05, \*\**P* < 0.01, ***P < 0.001, \*\*\*\**P* < 0.0001). **(D)** Addition of ROS scavenger *N*-acetylcysteine weakens the potentiation of benzydamine to doxycycline. **(E)** Benzydamine drastically impairs the function of multidrug efflux pumps in both MRSA T144 and *E. coli* B2. Rhodamine B (λexcitation = 540 nm, λemission = 625 nm) was used to characterize the activity of efflux pumps in bacteria. **(F)** Schematic illustrations of potentiating mechanisms of benzydamine with tetracyclines against drug-resistant pathogens.

### 2.5 Benzydamine restores doxycycline efficacy *in vivo* infection models

In view of the excellent synergy of doxycycline and benzydamine against MDR pathogens *in vitro*, we next tested whether they have potent activity in *vivo* with two animal infection models infected by MRSA T144 or *E. coli* B2. Firstly, we used a *G. mellonella* larvae infection model to explore their efficacy in *vivo*. As shown in **Figure 7A**, the infected larvae with the combination therapy of doxycycline plus benzydamine (50 + 50 mg/kg) resulted in above 80% survival during 5 days, which was higher than the doxycycline monotreatment (*P* = 0.0174 or 0.0397, corresponding to MRSA T144 and *E. coli* B2, respectively). In addition, the efficacy of this combination therapy in a neutropenic mouse thigh infection model was evaluated (**Figure 7B**). Similarly, the doxycycline plus benzydamine (50 + 10 or 50 + 50 mg/kg) significantly reduced bacterial burden in mice thighs compared with doxycycline alone (*P* < 0.0001). These data demonstrated the benzydamine plus doxycyline also has an excellent synergy effect *in vivo*.

**Figure 7.**
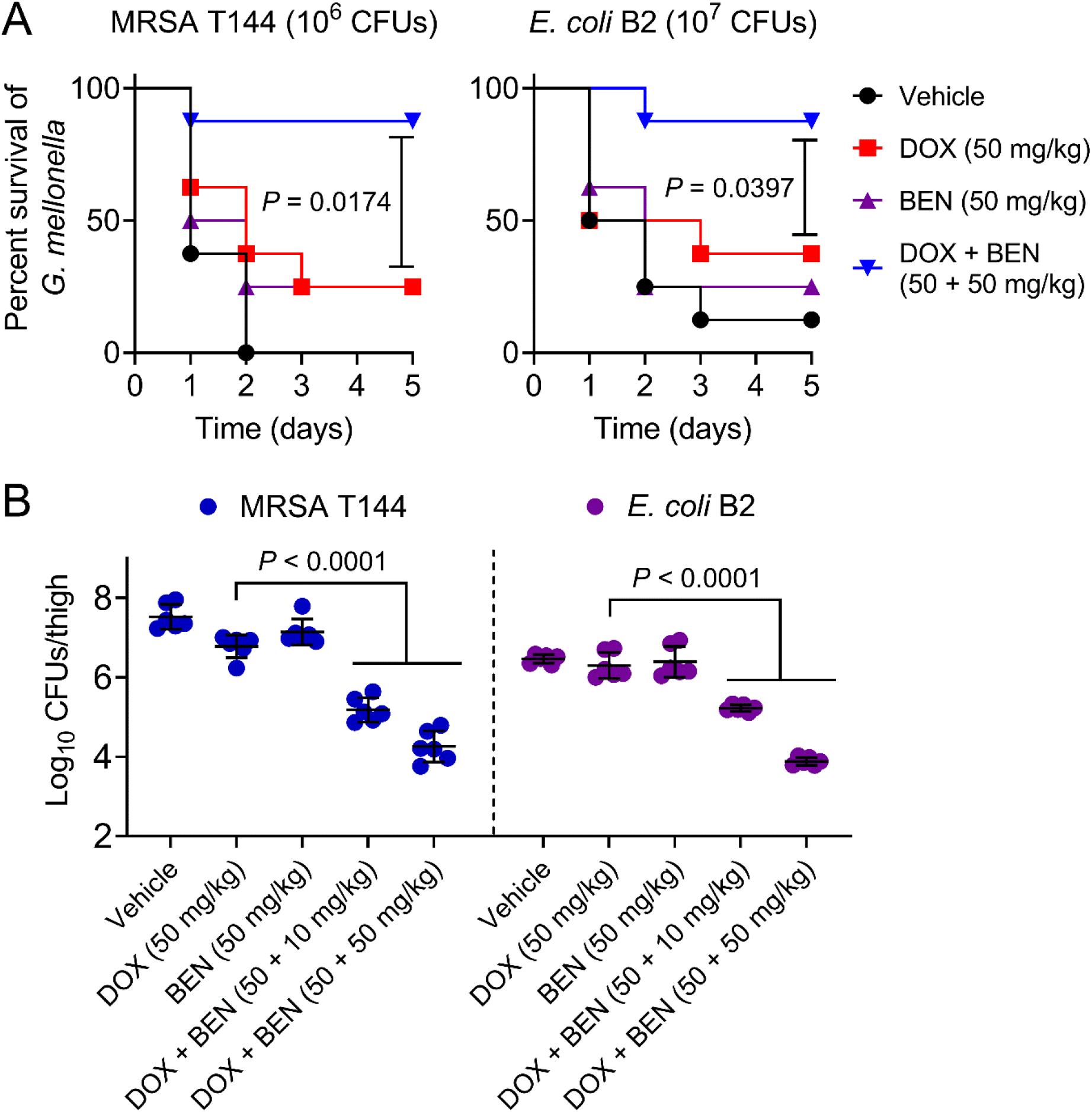
The combination of doxycycline and bezydamine is efficacious in various animals infection models. **(A)** Survival rates of *Galleria mellonella* (n = 8 per group) infected by MRSA T144 or *E. coli* B2 and then treated with doxycycline (50 mg/kg) or benzydamine (50mg/kg) alone or a combination of doxycycline plus benzydamine (50 + 50 mg/kg). **(B)** Combination of doxycycline (50 mg/kg) and benzydamine (50 mg/kg) significantly reduced the thigh bacterial loads of mice (n = 6 per group) infected by MRSA T144 or *E. coli* B2 (10^5^ CFUs per mouse) compared with doxycycline monotherapy (50 mg/kg).

## 3. Discussion

The development of multidrug resistance (MDR), extensive drug resistance (XDR), or pandrug resistance (PDR) phenotype in pathogenic bacteria undermines the clinical efficacy of antibiotics and leaves no available options for treatment of intractable bacterial infections (Falagas and Karageorgopoulos, 2008; Magiorakos et al., 2012). Despite many ongoing effects in identifying new classes of antimicrobial agents (Imai et al., 2019; Luther et al., 2019), few antibiotics have been approved for clinical use in the past 20 years. Accordingly, the average cost of research and development of a new drug from discovery to regulatory approval is US$2.6 billion, which takes more than 10 years, and the successful launch is less than one thousandth (Avorn, 2015; DiMasi et al., 2016). As such, alternative strategies are warranted to confront this serious global crisis. Repurposing previously approved non-antibacterial drugs as potential antibiotic adjuvants is gaining traction in both the public and private sector (Tyers and Wright, 2019). In this study, we revealed the adjuvant potential of benzydamine, a widely used non-steroidal anti-inflammatory drug in clinic, in combination with three classes of antibiotics against MDR *E. coli* B2. Most importantly, benzydamine displayed the greater synergistic activity with tetracyclines, which belong to broad-spectrum antibiotics. For various MDR pathogens, including MRSA, VRE, NDM/MCR-1/*tet*(X4)-expressing Gram-negative bacteria, the doxycycline-benzydamine combination showed unprecedented synergistic activity. Biofilm-producing bacteria are important causes of chronic and recurrent bacterial infection, but are commonly overlooked in the drug discovery. We found that the doxycycline-benzydamine combination is able to prevent the formation of *S. aureus* and *E. coli* biofilm, as well as eradicate the established biofilms. These data further support the notion that benzydamine is a great antibiotic adjuvant candidate to reverse bacterial resistance.

Bacterial energy metabolism such as proton motive force (PMF) plays a critical role in cellular activities including material transport, flagellar motility and ATP synthesis by the F_1_F_0_-ATPase (Paul et al., 2008). The disruption of PMF would inhibit the basic functions of bacteria and accelerate its death. Nevertheless, the PMF has remain largely unexplored as a target for the development of antimicrobial agents. Excitingly, using a deed learning approach, an new broad-spectrum antibiotic termed halicin was identified to selectively destroy the PMF (Stokes et al., 2020). Molecules I1-I3 and D1-D3, the potential modulators of PMF, showed killing activity against MRSA through preventing the electron transport and ATP synthesis (Farha et al., 2013). Generally, the PMF is comprised of two parameters: the electric potential (ΔΨ) and the transmembrane proton (ΔpH). Interestingly, to maintain the bacterial PMF, dissipation of either component would be compensatory increased by the another. In our study, we uncovered that benzydamine dissipated the ΔΨ component of the PMF in both Gram-positive and Gram-negative bacteria, in turn increased the ΔpH, which was critical for the uptake of tetracyclines. These findings was consistent with previous study that tetracyclines uptake is driven by ΔpH (Yamaguchi et al., 1991), whereas aminoglycosides uptake is highly dependent on ΔΨ (Taber et al., 1987). Toxicity concerns are critical factors that limit the clinical trials of new drugs (Segall and Barber, 2014). Meaningfully, in both *in vitro* and *in vivo* experiments, the combination of doxycycline and benzydamine exhibited negligible toxicity, indicating that the pre-clinical safety of this combination. Consistently, the successful paradigm of daptomycin depolarizing the cytoplasmic membrane provide a proof-of-concept for PMF-targeted antimicrobial agents (Hawkey, 2008). These findings suggest that bacterial PMF is a promising target for the development of novel antimicrobial agents and antibiotic adjuvants.

Furthermore, the doxycycline-benzydamine combination showed excellent synergistic bactericidal activity for all test MDR isolates, implying the actions of benzydamine is not merely for the promotion of tetracyclines uptake. Transcriptomic analysis coupled with phenotype experiments indicated that benzydamine not only triggered the generation of ROS, but also downregulated the GAD system that protects bacteria from oxidative damage. These modes of action resulted in the oxidative burst, which has been proved to be important for antibiotic-mediated killing. Additionally, the functions of MDR efflux pumps in bacteria were severely destroyed, in partly due to the dissipation of PMF by benzydamine. It would be interesting to investigate the potential of benzydamine as a new and broad-spectrum inhibitor of MDR efflux pumps.

To conclude, our findings revealed that non-steroidal anti-inflammatory drug benzydamine may serve as a novel antibiotic adjuvant to restore clinically relevant antibiotics activity particularly tetracyclines against infections caused by MDR pathogens. In addition, the identification and characterization of benzydamine demonstrate the remarkable potential of PMF downregulators as a feasible adjuvant therapy to tackle the escalating concern of antibiotic resistance.

## 4. Materials and methods

### Bacteria and reagents

All strains used in this study were listed in **Source data 1**. The bacteria were stored in nutrient broth supplemented with 20% (v/v) glycerol at −80°C. For experiments, all strains were grown in Mueller-Hinton Broth (MHB) or on LB agar (LBA) plates. Antibiotics were obtained from China Institute of Veterinary Drug Control and other chemical reagents were purchased from Aladdin (Shanghai, China) or TCI (Shanghai, China).

### MIC determinations

The MICs of all antibiotics and benzydamine were determined using broth dilution method, according to the CLSI 2018 guideline (In, 2018). All drugs were two-fold diluted in MHB and equally mixed with bacterial suspensions in a 96-well microtiter plate (Corning, New York, USA). After 16-20 h incubation at 37 °C, the MIC values were defined as the lowest concentrations of drugs with no visible growth of bacteria.

### Checkerboard analyses and FIC index determination

The fractional inhibitory concentrations (FIC) indices were measured by the checkerboard analyses (Song et al., 2020). Briefly, 100 μL of MHB was added into each well of a 96-well plate with 8×8 matrix, then the antibiotics and compounds were 2-fold diluted along the abscissa and ordinate, respectively. After incubated at 37 °C for 18 h with bacterial suspension (1.5 × 10^6^ CFUs/well), the optical density of each well at 600 nm were determined. The FIC was calculated as the MIC when the compound is used in combination divided by the MIC when it is used alone. The FIC index is the sum of the FICs of two compounds, and synergy is defined with FIC index ≤ 0.5.

### Hemolysis analysis

Hemolytic activity of doxycycline or in combination with benzydamine was assessed based on previous study (Liu et al., 2017). Briefly, 8% sheep blood cells was equal-volume incubated with 0 to 256 μg/mL doxycycline alone or in combination with 250 μg/mL benzydamine at 37 °C for 1 h. Phosphate buffer saline (PBS) and double-distilled water were used as negative and positive control, respectively. The absorption of released hemoglobin was measured at 576 nm by an Infinite M200 Microplate reader (Tecan, Männedorf, Switzerland). Hemolysis rate (%) was calculated by the result of absorbance of the sample subtracting the negative control divided by the positive control subtracting the negative control.

### *In vivo* toxicity of benzydamine-doxycycline combination

The *in vivo* toxicity was evaluated by gavaging a combination of doxycycline plus benzydamine (10 + 10 mg/kg) to female CD-1 mice (n = 6 per group). Mice were continuously gavaged for 6 days and body weights were recorded daily. On the seventh day, blood was collected for blood biochemical test and cell analysis.

### Inhibition of biofilm formation

MRSA T144 and *E. coli* B2 suspensions (1 × 10^7^ CFUs per mL) were exposed to doxycycline solutions (final concentrations ranging from 0.125 to 2 μg/mL) in the presence or absence of 50 μg/mL benzydamine. As negative control, bacteria were exposed to MHB without drugs. Bacteria were grown for 36 h at 37 °C under static conditions, and then 300 μL PBS was used to remove the planktonic bacteria. Then, added 200 μL of methanol to fix for 15 minutes, after that, the fixative were aspirated to air dry and 0.1% crystal violet was added for staining during 15 minutes. Dye solution was removed and stained-biofilm was washed three times with PBS and dried naturally. Lastly, crystal violet-stained biofilms were solubilized with 33% glacial acetic acid (100 μL) and incubated at 37 °C for 30 minutes. Biofilm mass was determined by monitoring the absorbance of supernatant at 570 nm (De et al., 2018).

### Biofilm eradication assay

Overnight MRSA T144 and *E. coli* B2 were diluted 1:100 into MHB and incubated at 37 °C with sharking at 200 rpm for 6 h. Subsequently, 100 μL bacterial suspensions were mixed with an equal volume of MHB in 96-well microtitre plate. After 36 h incubation at 37 °C, the planktonic bacteria were removed. Next, biofilms were treated with 32 to 256 μg/mL doxycycline alone or in combination with 50 μg/mL benzydamine. After 2 h incubation at 37 °C, the remaining cells was dispersed via ultrasonic treatment for 20 minutes. Finally, the mixed liquor was resuspended in sterile PBS and then the dilutions were plated on LBA plates and incubated overnight at 37 °C.

### Measurement of membrane potential

The membrane potential of *S. aureus* ATCC 29213, MRSA T144, *E. coli* ATCC 25922 and *E. coli* B2 was tested by the fluorescent probe DiSC_3_(5) (Aladdin, Shanghai, China). Bacterial cells were grown to the log phase in MHB, then washed with PBS to the OD_600_ of 0.5 and incubated with DiSC_3_(5) (0.5 × 10^−6^ M) for 30 min. Finally, varying concentrations of benzydamine (10 μL) were added into the 190 μL of DiSC_3_(5)-loaded cells. For all membrane potential experiments, the fluorescence intensity was measured with the excitation wavelength at 622 nm and emission wavelength at 670 nm using a Microplate reader (Tecan, Männedorf, Switzerland).

### Swimming motility experiment

0.3% (w/v) agar media composed of trypticase peptone (10 g/L), NaCl (10 g/L) and yeast extract (5 g/L) was used to assess bacterial swimming motility (Ejim et al., 2011). After the medium reached 50 °C, the final concentrations of benzydamine at 0, 31.25, 62.5, 125 and 250 μg/mL were added. A 2-µL volume of *S. aureus* 29213, MRSA T144, *E*.*coli* 25922 and *E*.*coli* B2 culture at an OD_600_ of 0.5 was placed in the center of each plate and allowed to stay for 30 min. The plates were placed in a 37 °C incubator for 48 h.

### Measurement of intracellular pH values

Overnight *S. aureus* ATCC 29213, MRSA T144, *E. coli* ATCC 25922 and *E. coli* B2 were resuspended to OD_600_ of 0.5 with PBS, and the final concentration of pH-sensitive fluorescent probe BCECF-AM (Ozkan and Mutharasan, 2002) (2 × 10^−6^ M for G^-^ bacteria and 0.5× 10^−6^ M for G^+^ bacteria) was added. After incubation for 30 min, four strains were treated with final concentration of benzydamine (125-1,000 μg/mL). The fluorescence intensity was immediately monitored with the excitation wavelength of 488 nm and emission wavelength of 535 nm.

### Uptake of doxycycline

Doxycycline uptake was evaluated by monitoring the fluorescence change of drug in bacteria (Ejim et al., 2011). Culture of MRSA T144 and *E. coli* B2 were grown to OD = 0.5. Cells were centrifuged at 3,500 rpm for 10 minutes and washed in the equal volume of PBS for three times. Subsequently, doxycycline at MIC alone or with final concentration of benzydamine at 250-1,000 μg/mL were added into the 96-wells plates containing cell suspensions at 100 µL/well. Infinite Microplate reader was used to monitor the fluorescence intensity with the excitation wavelength of 405 nm and emission wavelength of 535 nm.

### Measurement of ROS levels

The fluorescence probe 2′,7′-dichlorodihydrofluorescein diacetate (DCFH-DA, 10 μM) (Aranda et al., 2013) (Beyotime, Shanghai, China) was used to test the levels of ROS in *E. coli* B2 treated by benzydamine, doxycycline or their combination. After incubation 1 h, the fluorescence intensity was measured with the excitation wavelength of 488 nm and emission wavelength of 525 nm.

### Efflux pump assay

A fluorescence dye, rhodamine B (Forster et al., 2012) was applied to assay the inhibition of efflux pump of *E. coli* B2 and MRSA T144 treated by benzydamine. Bacterial cells were grown in MHB broth to mid-log phase (OD = 0.5) at 37 °C with shaking 200 rpm, then the cultures were washed and suspended with PBS. Subsequently, a final concentration of rhodamine B (Aladdin, Shanghai, China) (5 × 10^−6^ M) was added. After incubation at 37 °C for 30 min, probe-labeled cells were treated by doxycycline or benzydamine for 30 min, and the cultures washed and suspended with PBS containing 1% glucose. After incubation at 37 °C for 30 min, bacterial cells were centrifuged at 6,000 rpm at 4 °C for 5 min and supernatant was collected to determine with the excitation wavelength of 540 nm and emission wavelength of 625 nm.

### Time-dependent killing curves

Overnight culture of *E. coli* B2, *K. pneumoniae* D120 (*mcr-8*), *A. baumannii* C222 (*tet*(X6)), MRSA T144 and VRE A4 were diluted 1/1,000 in MHB, and incubated for 4 h at 37 °C with sharking at 200 rpm. Bacteria were then treated with either PBS, doxycycline (16 or 32 μg/mL) or benzydamine (250 μg/mL) alone or their combination. At the time points 0, 4, 8, 12, and 24 h, 100 μL aliquots were removed, centrifuged and resuspended in sterile PBS, the dilutions were plated on LBA plates and incubated overnight at 37 °C.

### Transcriptomic analysis

*E. coli* B2 were grown in MHB to the early exponential phase, then the final concentration of doxycycline (32 μg/mL) alone or in combination of benzydamine (250 μg/mL) was added. After incubation for 4 h, total RNA of culture was extracted by an EASYspin Plus kit (Aidlab, Beijing, China) and quantified by using a Nanodrop spectrophotometer (Thermo Scientific, MA, USA), and sequenced on Hiseq2000 with Truseq SBS Kit v3-HS (200 cycles) (Illumina) with the read length as 2 × 100 (PE100). Raw sequencing reads were filtrated and mapped against the reference genome of *E. coli* K-12. The FPKM (Fragments Per Kilobase of transcript per Million mapped reads) method was used to identify differentially-expressed genes with p-values ≤ 0.05 and fold change (FC) values ≥ 2 (log2 FC ≥ 1 or log2 FC ≤ -1). Differences between these two treatments were studied by Cuffdiff program (http://cufflinks.cbcb.umd.edu/)

### *Galleria mellonella* infection model

*Galleria mellonella* larvae (Huiyude Biotech Company, Tianjin, China) were divided into four groups (n = 8 per group) and infected with MRSA T144 (10^6^ CFU_S_) or *E. coli* B2 (10^7^ CFU_S_) suspension. After 1 h post infection, group 1 was subjected to PBS treatment, groups 2 and 3 were treated with doxycycline or benzydamine (50mg/kg) respectively, group 4 was treated with doxycycline plus benzydamine (50 + 50 mg/kg). Survival rates of *Galleria mellonella* larvae were recorded for 5 days.

### Neutropenic mouse thigh infection model

6-8-week-old female CD-1 mice were obtained from Comparative Medicine Centre of Yangzhou University (Jiangsu, China). Mice studies were performed in accordance with the guidelines of Jiangsu Laboratory Animal Welfare and Ethical of Jiangsu Administrative Committee of Laboratory Animals. The protocols for all animal studies were approved by Jiangsu Administrative Committee for Laboratory Animals (Permission number: SYXKSU-2007-0005). The laboratory animal usage license number is SCXK-2017-0044, certified by Jiangsu Association for Science and Technology.

Female CD-1 mice (n = 6 per group) were firstly treated by cyclophosphamide with 150 mg/kg in the 4 days before infection, and 100 mg/kg in the 1 day before infection. MRSA T144 or *E. coli* B2 suspension (100 μL, 10^5^ CFUs per mouse) was injected into the right thighs of mice. After 2 h post infection, mice were intraperitoneally injected with PBS, doxycycline (50 mg/kg), benzydamine (50 mg/kg), or combinations (50 + 10 mg/kg, 50 + 50 mg/kg). At 48 h post infection, mice were euthanized by cervical dislocation. The right thigh muscle was aseptically removed, homogenized, serially diluted and plated on LBA to count bacterial numbers.

## Statistical analyses

Statistical analysis was performed using GraphPad Prism version 8.3.0. All data was shown as mean ± SD. Unpaired *t*-test between two groups or one-way ANOVA among multiple groups were used to calculate *P*-values (\**P* < 0.05, \*\**P* < 0.01, \*\*\**P* < 0.001).

## Acknowledgments

This work was supported by the National Natural Science Foundation of China (32002331), National Key Research and Development Program of China (2018YFA0903400), Natural Science Foundation of Jiangsu Province of China (BK20190893), Agricultural Science and Technology Independent Innovation Fund of Jiangsu Province (CX(20)3091), China Postdoctoral Science Foundation funded project (2019M651984), A Project Funded by the Priority Academic Program Development of Jiangsu Higher Education Institutions (PAPD) and Lift Engineering of Young Talents of Jiangsu Association for Science and Technology.

## Author contributions

Z.W. and Y.L. design and conceived the project. Z.T., J. S., T. D. and Y.J. performed all experiments. Y.L., Z.T., R.L. and X.X. analyzed the data. Y.L. and Z. T. wrote the manuscript. All authors read and approved the manuscript.

## Additional files

- **Source data 1**. Bacterial strains used in this study.
- **Source data 2**. Differentially expressed genes (DEGs) of *E. coli* B2 treated with doxycycline plus benzydamine.
- **Transparent reporting form**

---

**This article includes the following four figure supplements for figure 1:**

**Figure 1-figure supplement 1.**
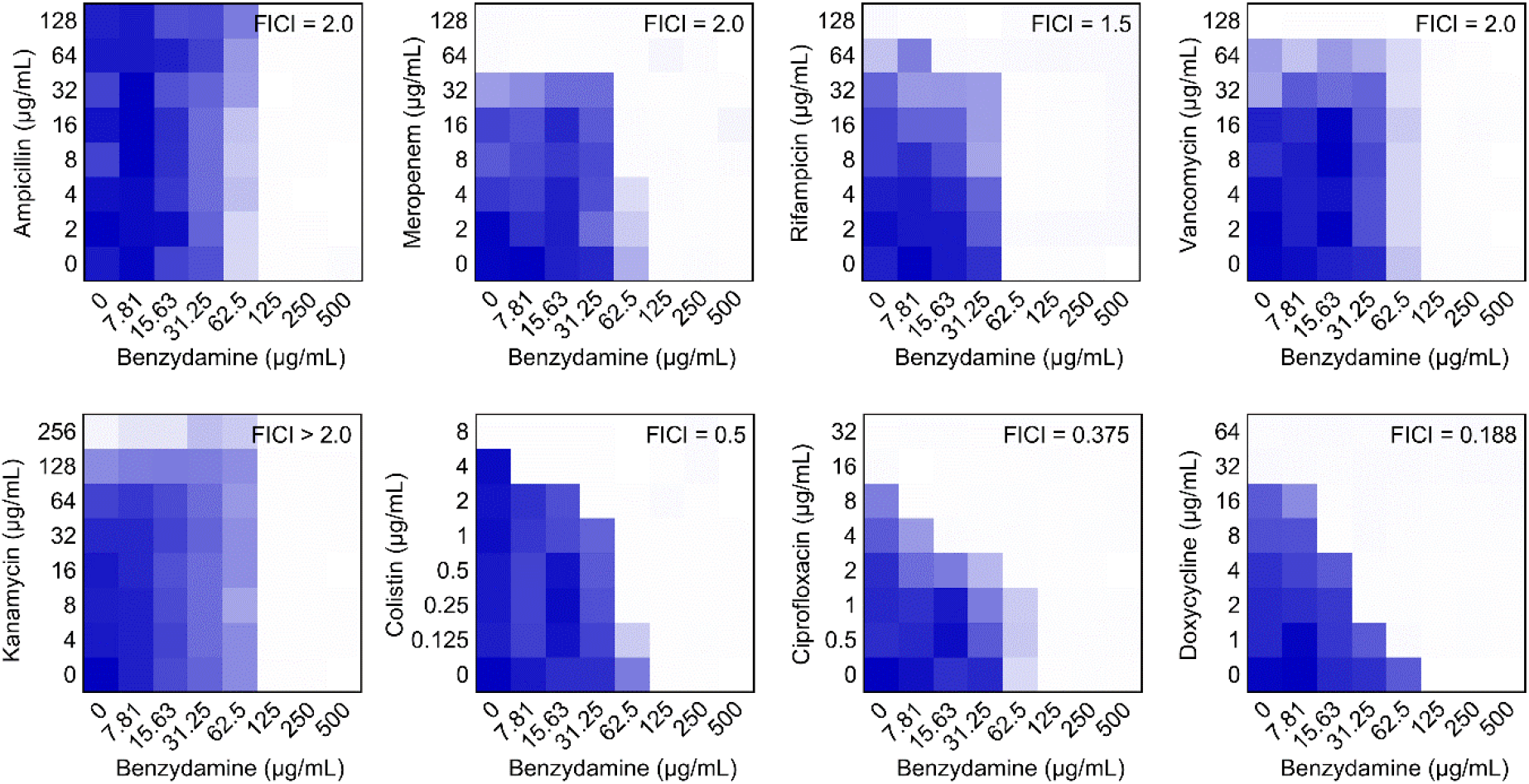
Interaction between benzydamine and multiple classes of antibiotics against *E. coli* B2 by checkerboard assay, related to Table S2. Dark blue regions represent higher cell density. Data represent the mean OD (600 nm) of two biological replicates.

**Figure 1-figure supplement 2.**
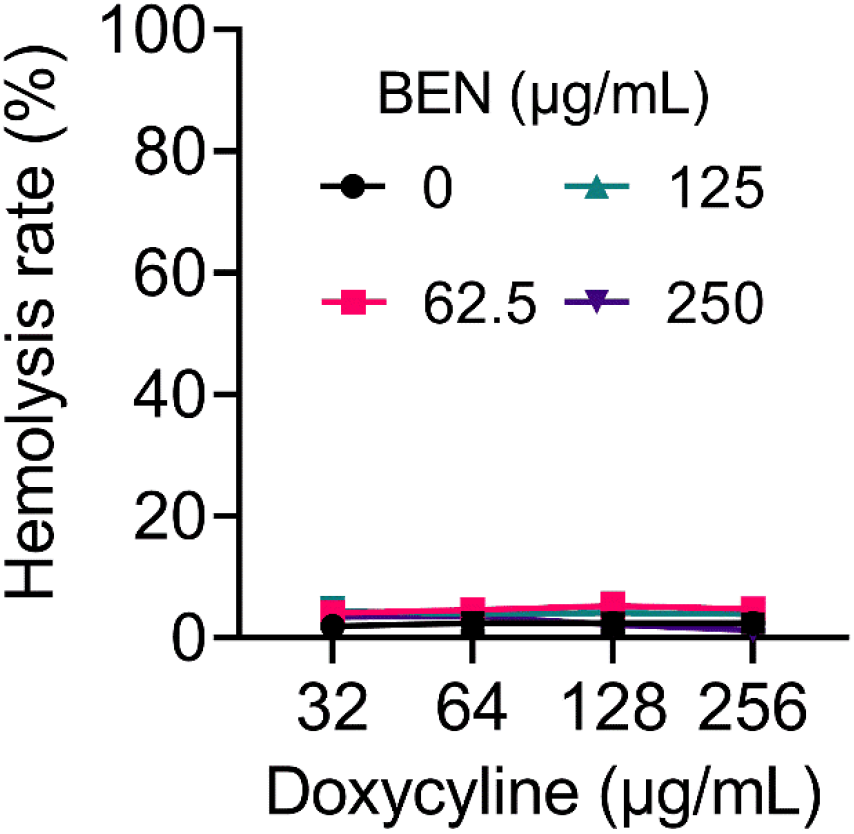
Doxycycline plus benzydamine displays negligible hemolytic activity on mammals’ RBCs. Phosphate buffer saline (PBS) and double-distilled water were used as negative and positive control, respectively.

**Figure 1-figure supplement 3.**
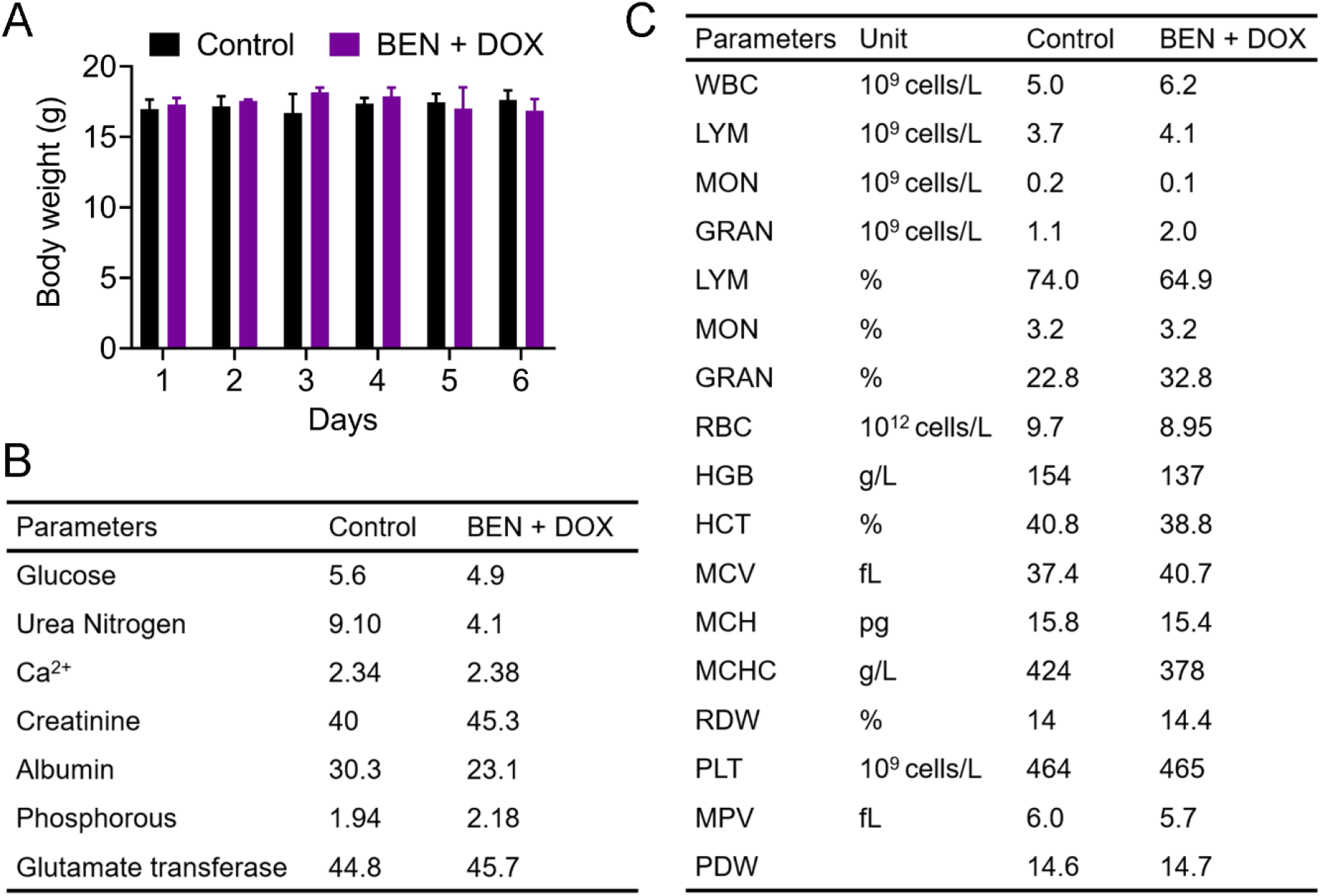
*In vivo* toxicity evaluation of the combination of benzydamine and doxycycline. CD-1 female mice (n = 6 per group) were gavaged with vehicle or the bezydamine-doxycycline combination once daily for six days. Meanwhile, the mice body weight (A), serum biochemical analysis (B) and whole-blood cell analysis (C) were shown. The data were presented as mean. White blood cell (WBC), lymphocyte (LYM), monocyte (MON), neutrophils (NEU), red blood cell (RBC), hemoglobin (HGB), hematocrit (HCT = RBC%), the mean corpuscular volume (MCV, average volume of red cells), mean corpuscular hemoglobin (MCH), platelet count (PLT), and mean corpuscular hemoglobin concentration (MCHC, the average amount of hemoglobin inside a single red blood cell).

**Figure 1-figure supplement 4.**
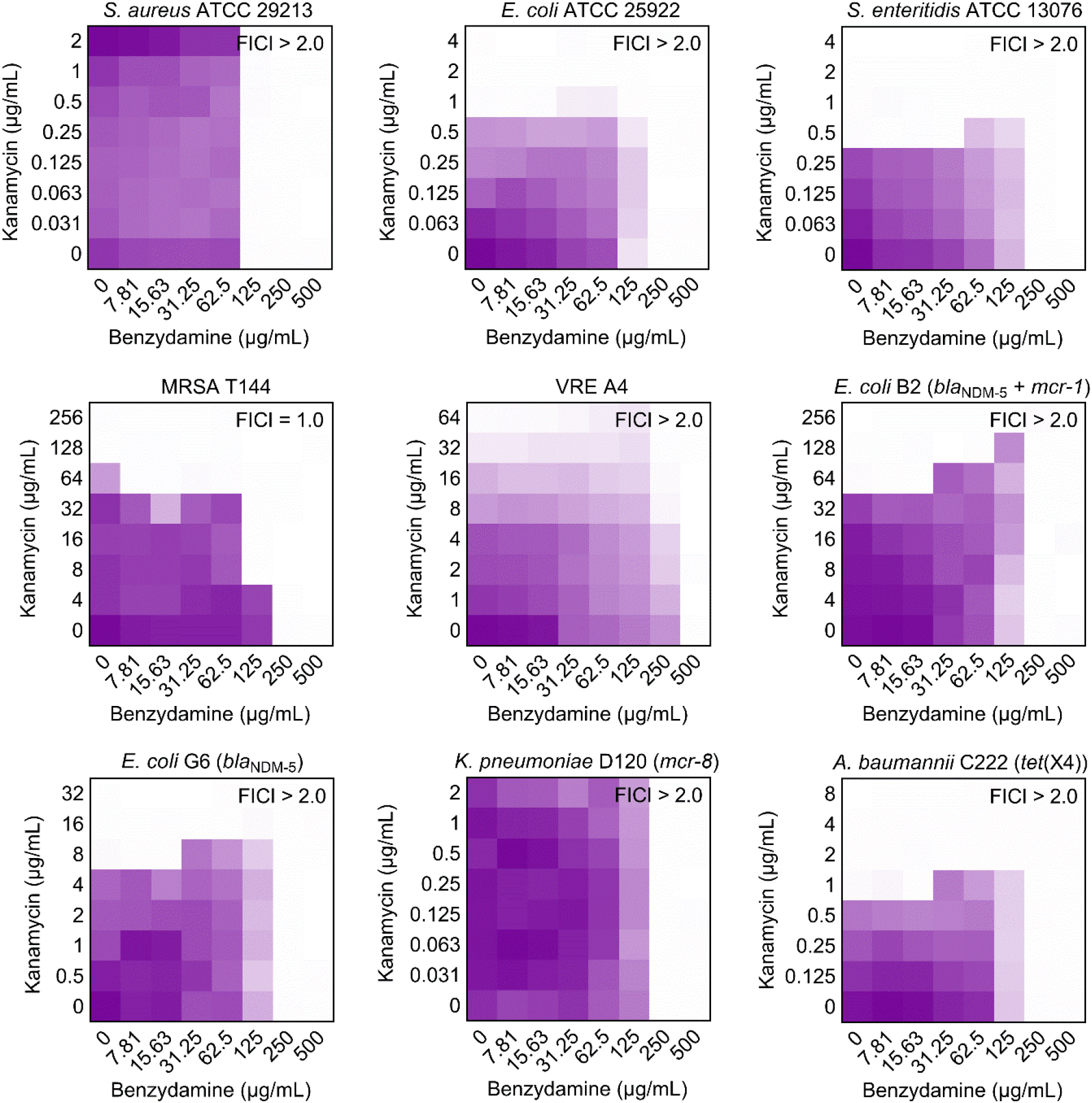
Antagonism effect of benzydamine in combination with kanamycin in both kanamycin-sensitive and resistant bacteria. Dark blue represents greater growth. Data represent the mean OD (600 nm) of two biological replicates.

**This article includes the following one figure supplement for figure 4:**

**Figure 4-figure supplement 1.**
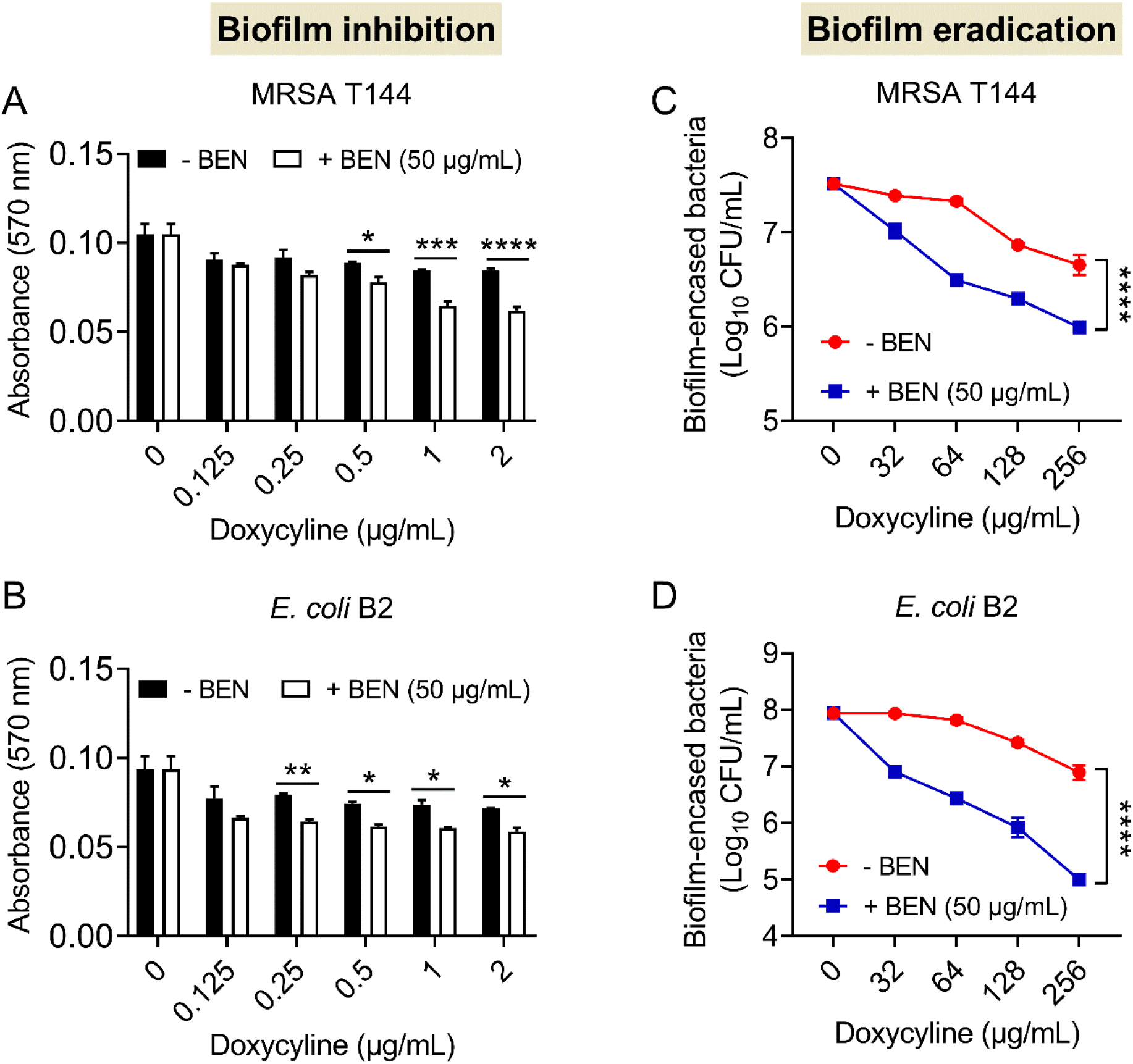
Benzydamine enhances the biofilm inhibition and eradication activities of doxycycline. **(A and B)** Benzydamine supplementation potentiates the inhibitory effect of doxycycline on MRSA T144 (A) and *E. coli* B2 (B) biofilm formation. **(C and D)** Addition of benzydamine drastically promotes the eradication of established biofilm of MRSA T144 (A) and *E. coli* B2 (B) by doxycycline.

**This article includes the following one figure supplement for figure 5:**

**Figure 5-figure supplement 1.**
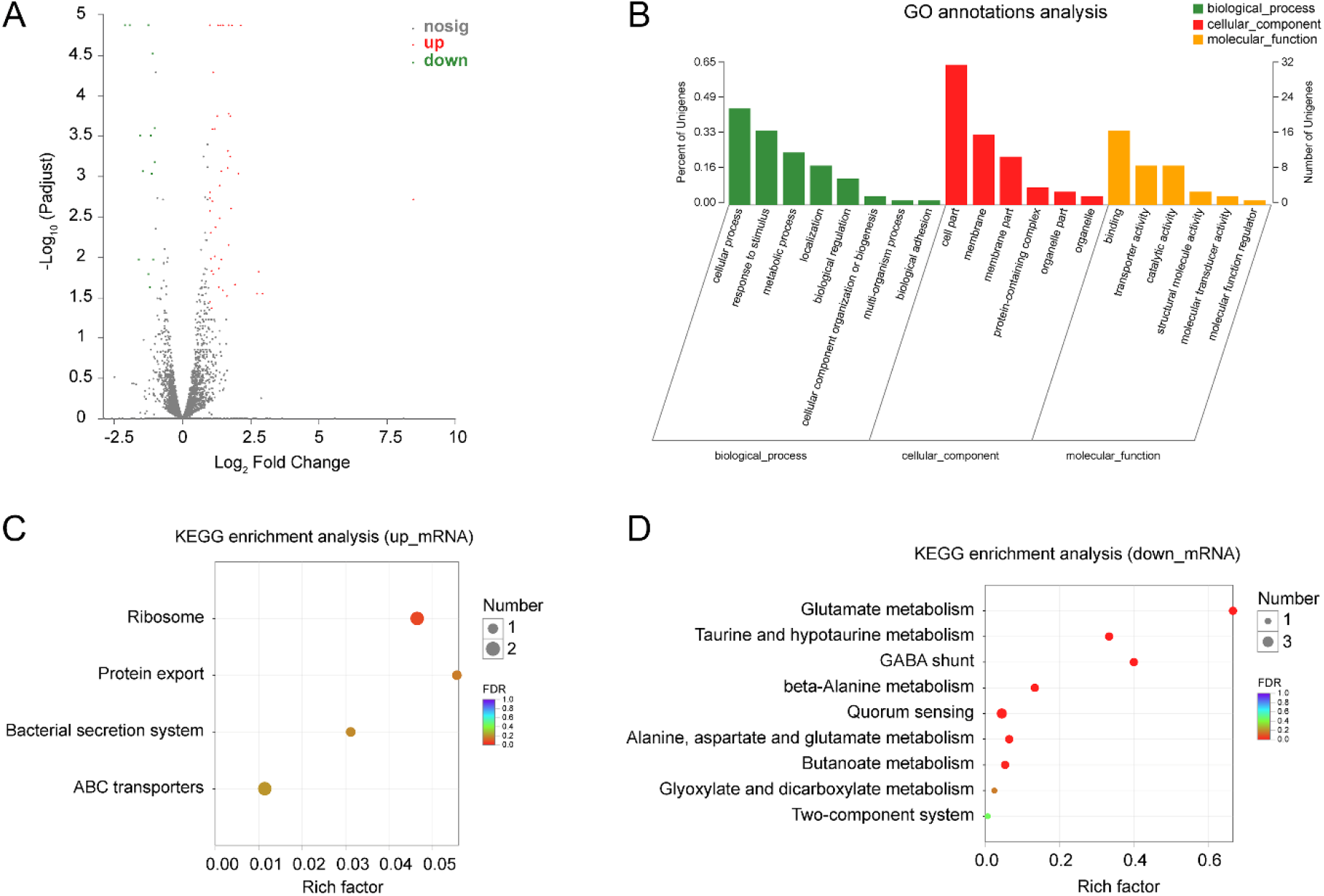
Transcriptomic analysis of *E. coli* B2 after exposure to doxycycline or in combination with benzydamine. **(A)** Volcano plot and **(B)** GO (gene ontology) annotation analysis of the differential expression genes (DEGs) in *E. coli* B2 after exposing doxycycline (32 µg/mL) or the combination of doxycycline (32 µg/mL) plus benzydamine (250 µg/mL) for 4 h.The x- and y-axes in (A) represent the expression changes and corresponding statistically significant degree, respectively. An adjusted p-value < 0.05 (Student’s *t*-test with Benjamini–Hochberg false discovery rate adjustment) and |log2 Fold change| ≥1 were applied as the cutofffor significant DEGs. KEGG (Kyoto Encyclopedia of Genes and Genomes) enrichment analysis of **(C)** upregulated DEGs and **(D)** downregulated DEGs. The 10 most significant enriched pathways are shown.

